# Context-dependent variant interpretation from Mendelian disease to genetic predisposition: a proof-of-concept using *LPL*

**DOI:** 10.64898/2026.08.12.744351

**Authors:** Qi Yang, Wen-Bin Zou, Na Pu, Yi Li, Yuepeng Hu, Yuan-Chen Wang, Xin Liu, Emmanuelle Génin, Emmanuelle Masson, Jiucun Wang, Claude Férec, David N. Cooper, Weiqin Li, Jian-Min Chen

## Abstract

As genomic sequencing evolves beyond rare disease diagnostics toward population screening and precision medicine, clinical variant interpretation is increasingly challenged by variants whose clinical consequences depend on biological context. Current frameworks, including the ACMG/AMP guidelines, generally assign a single classification to each variant regardless of inheritance state or genetic context, potentially failing to communicate context-dependent clinical consequences. Here, we address this issue using loss-of-function variants in *LPL* as a uniquely informative model system in which residual physiological LPL activity can be directly quantified *in vivo*. By systematically integrating published biallelic *LPL* genotypes, physiological measurements, functional studies, and clinical phenotypes, we identified a biologically meaningful transition at approximately 10% residual physiological LPL activity. Activity below this level was predominantly associated with classical childhood-onset familial chylomicronemia syndrome (FCS), whereas higher activity was associated with phenotypic attenuation and modifier-dependent clinical expression. Furthermore, heterozygous loss-of-function variants exhibited an estimated penetrance of 5–7% for severe hypertriglyceridemia. We therefore propose a context-dependent framework in which biallelic complete- or near-complete loss-of-function genotypes are interpreted as causative for FCS, whereas heterozygous variants are interpreted as predisposing to severe hypertriglyceridemia while retaining recognition of FCS carrier status. Together, our findings demonstrate that clinical variant interpretation should integrate available biological context—including, where relevant, allelic configuration, residual biological function, and penetrance—rather than rely on the intrinsic molecular consequence of the variant alone. More broadly, this framework provides a conceptual model for interpreting variants across the continuum from Mendelian disease to genetic predisposition in the era of precision medicine.

## Introduction

The widespread implementation of exome and genome sequencing has transformed medical genetics from a field focused primarily on diagnosing rare Mendelian disorders into one supporting population screening, disease prevention, and precision medicine [1–4]. As a result, clinical variant interpretation now extends far beyond determining whether a variant causes a classical Mendelian disease. In many settings, the underlying genetic architecture becomes apparent only after molecular analysis, and even disorders initially considered to follow simple Mendelian inheritance may instead exhibit incomplete penetrance, oligogenic inheritance, or gene–environment interactions. This broader application of genomic sequencing has exposed important limitations of current clinical variant interpretation frameworks, including the American College of Medical Genetics and Genomics and Association for Molecular Pathology (ACMG/AMP) guidelines [5], which were originally developed to support the molecular diagnosis of classical Mendelian disorders.

Several conceptual and methodological advances have sought to improve clinical variant interpretation [6–13]. Among these, we previously proposed expanding the ACMG/AMP framework by incorporating a predisposing variant category for variants that reproducibly increase disease susceptibility without exhibiting the high penetrance expected of pathogenic variants causing classical Mendelian disease [7, 13]. Beyond this proposed expansion, we emphasized that clinical variant interpretation should be performed within the specific gene–disease context because functional tolerance thresholds and modes of inheritance vary among gene–disease systems [13].

Despite these advances, current frameworks generally assign a single classification to each variant even when the same variant has fundamentally different clinical consequences in different biological contexts. This creates a particularly important challenge for variants in genes traditionally associated with autosomal recessive disease. In these systems, heterozygous carriers have classically been regarded as clinically unaffected. Yet, accumulating evidence demonstrates that variants causing recessive Mendelian disease in the biallelic state may also confer susceptibility to complex or multifactorial disease in the heterozygous state [14, 15]. Such context-dependent clinical consequences remain incompletely captured in current clinical databases and the literature, potentially hampering the full realization of precision medicine [16, 17].

This challenge is exemplified by *LPL* (MIM: 609708), which encodes lipoprotein lipase, the principal enzyme responsible for plasma triglyceride hydrolysis. Biallelic complete- or near-complete loss-of-function (c/nc-LoF) variants in *LPL* account for approximately 80–90% of cases of familial chylomicronemia syndrome (FCS), a rare autosomal recessive disorder characterized by severe refractory hypertriglyceridemia (HTG), recurrent pancreatitis, and profound physiological LPL deficiency [18, 19]. By contrast, heterozygous c/nc-LoF *LPL* variants are well-established genetic predisposing factors for severe HTG [20–22], which itself is associated with an increased risk of acute pancreatitis [23]. Despite these distinct clinical consequences, c/nc-LoF *LPL* variants, irrespective of zygosity, are frequently classified as pathogenic in ClinVar (https://www.ncbi.nlm.nih.gov/clinvar/; last accessed 7 July 2026) and the literature [20, 21, 24, 25] through extrapolation from their established role in FCS.

Resolving this challenge requires moving beyond a static, one-variant–one-label paradigm. Here, we use *LPL* as a proof-of-concept model to establish a biologically grounded, context-dependent framework for variant interpretation across the continuum from Mendelian disease to genetic predisposition. Unlike most disease-associated genes, *LPL* provides a uniquely informative model system because residual physiological enzyme activity can be quantified directly *in vivo* through post-heparin plasma LPL activity measurements. This unique feature establishes a quantitative bridge between molecular variant effect, residual physiological function, and clinical consequence, making *LPL* an ideal system for investigating context-dependent variant interpretation. By systematically integrating published human *LPL* genotypes, residual physiological LPL activity, functional studies, clinical phenotypes, and penetrance estimates for heterozygous c/nc-LoF variants, we identify reproducible biological reference points linking different levels of residual physiological function to distinct clinical consequences. Together, these analyses establish a biologically grounded framework for context-dependent variant interpretation across the continuum from Mendelian disease to genetic predisposition.

## Results

### Assembly and classification of published biallelic *LPL* genotypes

To establish a biological framework linking residual physiological LPL activity to clinical consequence, we focused on the functional spectrum spanning complete physiological LPL deficiency to the theoretical 50% residual activity expected for heterozygous complete-LoF variants. This spectrum captures the transition from classical Mendelian FCS to genetic predisposition for severe HTG and provides the basis for the subsequent analyses.

To investigate how residual physiological LPL activity relates to clinical consequence, we assembled a curated dataset of published human biallelic *LPL* genotypes for which both physiological (*in vivo*) and functional (*in vitro*) LPL activity measurements were available. This dataset enabled direct comparison of residual physiological enzyme activity with clinical phenotype across a broad spectrum of residual LPL function. Applying predefined inclusion criteria yielded 45 published biallelic *LPL* genotypes containing at least one missense variant for which both *in vivo* and *in vitro* LPL activity had been reported in the same study (Fig. 1).

**Figure 1.**
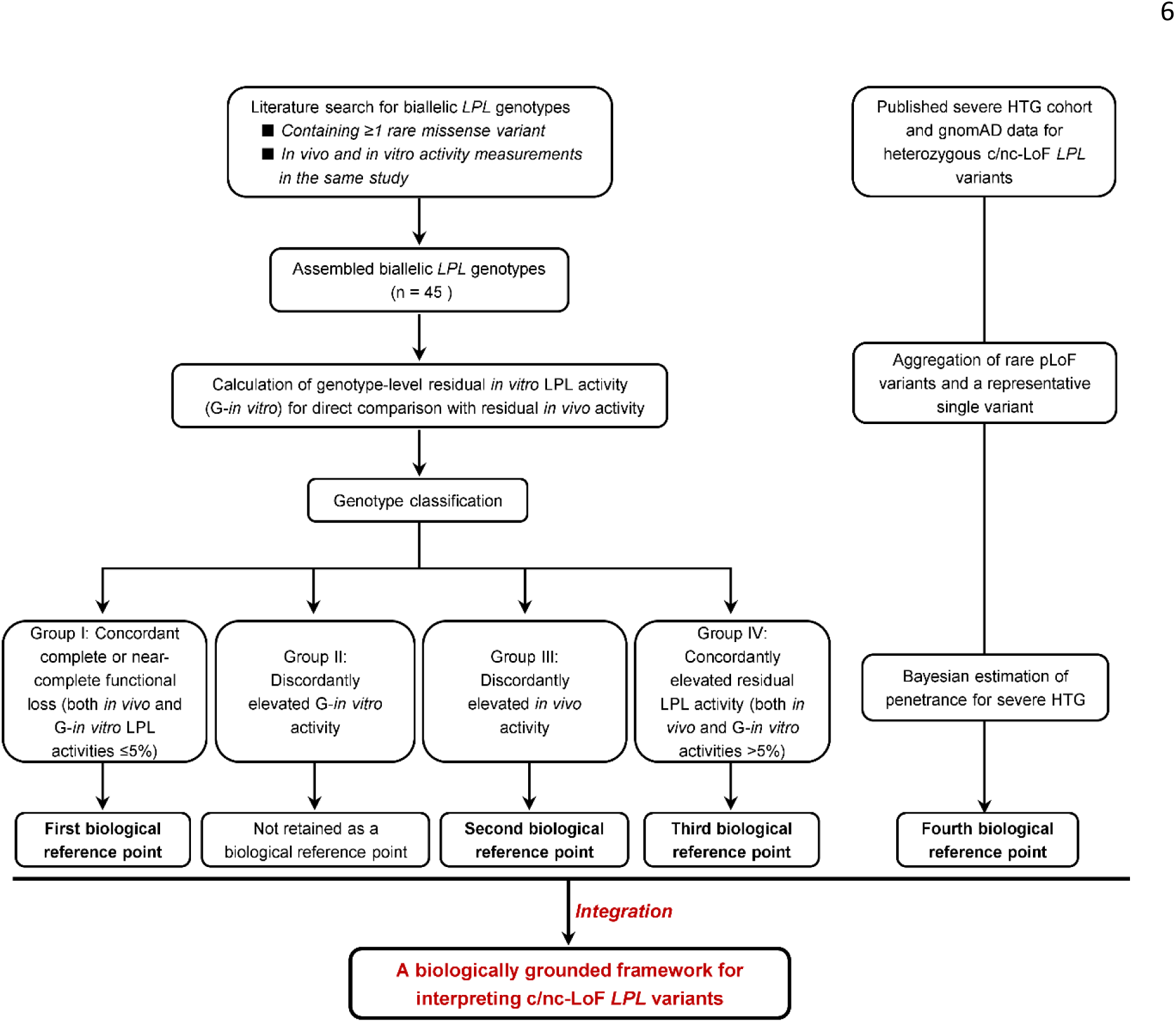
Overall analytical strategy. The study combined two complementary analyses. Published biallelic *LPL* genotypes were used to define biological reference points linking residual physiological LPL activity to clinical consequences, whereas Bayesian penetrance analysis of heterozygous c/nc-LoF variants established the fourth biological reference point for variant classification. Integration of these complementary lines of evidence yielded a biologically grounded framework for interpreting c/nc-LoF *LPL* variants. *Abbreviations:* c/nc-LoF, complete- and near-complete loss-of-function; HTG, hypertriglyceridemia; pLoF, predicted loss-of-function.

Because the dataset was derived from studies published over several decades, methodological heterogeneity across studies—and, in some instances, within individual studies—was unavoidable. To account for this heterogeneity, we systematically examined the relationships among residual *in vivo* LPL activity, genotype-level *in vitro* (G-*in vitro*) LPL activity, and clinical phenotype. To facilitate this analysis, a threshold of 5% of normal activity was used to initially classify genotypes according to the concordance between residual *in vivo* and G-*in vitro* activity. Genotypes showing concordant complete or near-complete functional loss (*in vivo* and G-*in vitro* ≤5%) and those showing concordantly elevated residual activity (*in vivo* and G-*in vitro* >5%) constituted the two concordant groups. The remaining genotypes showed either discordantly elevated *in vivo* activity or discordantly elevated G-*in vitro* activity and constituted the two discordant groups. These four groups provided an initial analytical framework, with the relationships among residual *in vivo* activity, G-*in vitro* activity, and clinical phenotype subsequently examined in detail within each group.

### Group I: Concordant complete or near-complete functional loss

Group I comprised 27 biallelic genotypes in which both residual *in vivo* and G-*in vitro* LPL activities were ≤5% of normal (Supplementary Table S1), making it the largest group in the dataset.

The clinical manifestations associated with Group I were highly consistent with FCS. All 27 corresponding individuals were diagnosed with FCS, LPL deficiency, type I hyperlipoproteinemia, or severe primary HTG. Because classical FCS typically presents during infancy or childhood, age at symptom onset—or, when unavailable, age at genetic diagnosis—was extracted whenever possible to assess early disease presentation (≤10 years). Early presentation was documented in 23 of the 27 individuals (85%). For the remaining four individuals, genetic diagnosis occurred between 18 and 39 years of age, although age at symptom onset was not reported (Supplementary Table S1).

Taken together, the concordant complete or near-complete functional deficiency indicated by both residual *in vivo* and G-*in vitro* LPL activity, together with the highly consistent phenotype of classical FCS, establishes Group I as the first biological reference point in our analytical framework. This reference point anchors the highly penetrant Mendelian disease end of the biological continuum linking residual physiological LPL activity to clinical consequence.

### Group II: Discordantly elevated G-*in vitro* activity

Only three biallelic genotypes were classified as Group II, exhibiting discordantly elevated G-*in vitro* activity (15.3–35% of wild-type) despite residual *in vivo* LPL activity of <1% (Supplementary Table S2) [26–28].

The first two genotypes (c.382A>G (p.Thr128Ala)/c.829G>A (p.Asp277Asn) [26] and c.662T>C (p.Ile221Thr)/c.1334G>A (p.Cys445Tyr) [27]) nevertheless exhibited clinical phenotypes consistent with classical early-onset FCS (Supplementary Table S2). Despite G-*in vitro* activities of 15.3% and 25% of wild-type, residual *in vivo* LPL activity was virtually absent (<0.9% and 0%, respectively). These findings suggest that the corresponding *in vitro* functional measurements may not have accurately reflected residual physiological LPL function *in vivo*, as noted by one of the original studies [27].

By contrast, the third genotype (c.953A>G (p.Asn318Ser)/c.1019-3C>A) was associated with an atypical later-onset phenotype. Although the original study reported no detectable *in vitro* activity for p.Asn318Ser [28], several subsequent studies demonstrated substantial residual activity (approximately 50–70% of wild-type) [29–32]. Consequently, the G-*in vitro* activity used in the present analysis was derived from these later studies and is more consistent with the atypical clinical presentation observed in this case (Supplementary Table S2).

Taken together, Group II does not provide a sufficiently robust basis for establishing a biological reference point linking residual physiological LPL activity to clinical consequence.

### Group III: Discordantly elevated *in vivo* LPL activity

Group III initially comprised 11 biallelic genotypes with residual *in vivo* LPL activity ranging from approximately 6% to 27.6% of normal despite G-*in vitro* activity within the complete or near-complete LoF range.

Three individuals representing two homozygous genotypes (c.693C>G (p.Asp231Glu), n = 2; c.809G>A (p.Arg270His), n = 1) exhibited the highest reported residual *in vivo* activities (17%– 27.6%) despite complete absence of G-*in vitro* activity (Supplementary Table S3). As discussed in Supplementary Note 1, these observations most likely reflect case-specific methodological and/or biological factors. Consequently, subsequent analyses focused on the remaining nine Group III genotypes.

Among the remaining nine genotypes (Table 1) [26, 33–38], three overall patterns emerged. First, residual *in vivo* LPL activity was consistently modest, ranging from approximately 6% to 10% of normal. Second, childhood presentation was less frequent than in Group I. Among individuals with available age information, presentation at ≤10 years of age was observed in 23 of 27 (85%) Group I individuals but only 4 of 9 (44%) Group III individuals. Third, within the complete or near-complete LoF range, detectable residual G-*in vitro* activity was more common than in Group I. Complete absence of G-*in vitro* activity was observed in 21 of 27 (78%) Group I genotypes compared with only 2 of 9 (22%) Group III genotypes.

**Table 1.** Group III biallelic *LPL* genotypes with discordantly elevated *in vivo* LPL activity.

| Biallelic genotype | Reference | Zygosity | Post-heparin plasma LPL (% of controls) | Reason for genetic analysis | Age at genetic analysis | Disease history | *In vitro* LPL activity (% of WT activity)<sup>a</sup> |
| --- | --- | --- | --- | --- | --- | --- | --- |
| c.3G>C (p.Met1?) / c.[3G>C;805G>A] (p.[Met1?;Glu269Lys]) | Yu et al. [33] | Compound | 6.5 | Acute pancreatitis and severe HTG | NR | Hospitalized at age 18 with acute pancreatitis, when severe HTG was discovered. LPL deficiency was subsequently diagnosed based on markedly reduced post-heparin plasma LPL activity. | p.Met1?: 1.7<br>p.Glu269Lys: 0<br>p.[Met1?;Glu269Lys]: 0<br>*G-in vitro*: 0.85 |
| c.300C>A (p.Tyr100*) / c.306A>C (p.Arg102Ser) | Wilson et al. [34] | Compound | ~6 | Type I hyperlipoproteinemia | 43 y | Lipemia was noted in infancy. The patient recalled recurrent episodes of abdominal pain precipitated by dietary fat throughout childhood. | p.Tyr100*: 0<br>p.Arg102Ser: <1<br>*G-in vitro*: <0.5 |
| c.506G>A (p.Gly169Glu) | Ameis et al. [35] | Homozygous | 8.3 | Type I hyperlipoproteinemia | 22 y | Presented at 27 days of age with plasma triglyceride level of 337 mmol/L, lipemia retinalis, and eruptive xanthomas. | *G-in vitro*: 0 |
| c.553G>A (p.Ala185Thr) | Mailly et al. [36] | Homozygous | Described as low (<10) | LPL deficiency | 6 m | Severe HTG, fasting chylomicronemia, and hepatosplenomegaly | *G-in vitro*: 3.2 |
| c.643G>A (p.Gly215Arg) | Benlian et al. [26] | Homozygous | 8 | FCS | 52 y | Diagnosed at age 29 years (triglyceride, 42.9 mmol/L). The patient reported abdominal pain induced by high dietary fat intake but had no history of acute pancreatitis. | *G-in vitro*: <1 |
| c.658A>C (p.Ser220Arg) / 2 kb insertion | Mailly et al. [36] | Compound | Described as low (<10) | Type V lipid profile | 25 y | Abdominal pain and severe HTG | p.Ser220Arg: 2.0<br>2 kb insertion: 0<br>*G-in vitro*: 1.0 |
| c.809G>A (p.Arg270His) / c.829G>A (p.Asp277Asn) | Ishimura-Oka et al. [37] | Compound | 6.4<sup>b</sup> | Type I hyperlipoproteinemia | 15 y | Presented with recurrent vomiting at 3 months of age and was found to have hyperlipidemia. Diagnosed with type I hyperlipoproteinemia at 7 years of age. Lipemia retinalis and splenomegaly were reported. | p.Arg270His: 0<br>p.Asp277Asn: 3<br>*G-in vitro*: 1.5 |
| c.909G>C (p.Leu303Phe) | Saika et al. [38] | Homozygous | 6 | Type I hyperlipoproteinemia | 55 y | First hospitalized at age 23 years because of acute pancreatitis, at which time markedly elevated plasma triglyceride levels were documented. | *G-in vitro*: 0 |
| c.938T>C (p.Leu313Pro) / c.290_293delinsGG (p.Ala97Glyfs*50) | Benlian et al. [26] | Compound | 6.6 | FCS | 67 y | History of abdominal pain induced by high dietary fat intake and hyperlipidemia. At age 53 years, an episode of acute pancreatitis was accompanied by eruptive xanthomas and chylomicronemia (triglyceride, 144.5 mmol/L). | p.Leu313Pro: <1<br>p.Ala97Glyfs*50: 0<br>*G-in vitro*: <0.5 |

<sup>a</sup>*G-in vitro* denotes the genotype-level estimate of *in vitro* LPL activity derived from the activities of the constituent variant(s) (see Methods).
<sup>b</sup>The range of post-heparin plasma LPL activity of normal subjects was 5-20 $\mu\text{mol/h}$ per ml [37]. For calculation, a normal value of 12.5 $\mu\text{mol/h}$ per mL was used.
*Abbreviations*: HTG, hypertriglyceridemia; LPL, lipoprotein lipase; y, years.

Taken together, Group III identifies residual physiological LPL activity of approximately 6–10% as a potentially informative range associated with heterogeneous clinical expression. The biological significance of this activity range is further evaluated together with the findings from Group IV in the following section.

### Group IV: Concordantly elevated residual LPL activity

Group IV comprised four biallelic genotypes characterized by both residual *in vivo* and G-*in vitro* LPL activities >5%, although the magnitudes of the two measures were not closely correlated within individual genotypes (Table 2) [39–42]. Specifically, residual *in vivo* activities ranged from 12% to 40% of normal, whereas G-*in vitro* activities ranged from 5.2% to 20% of wild-type activity.

**Table 2.**
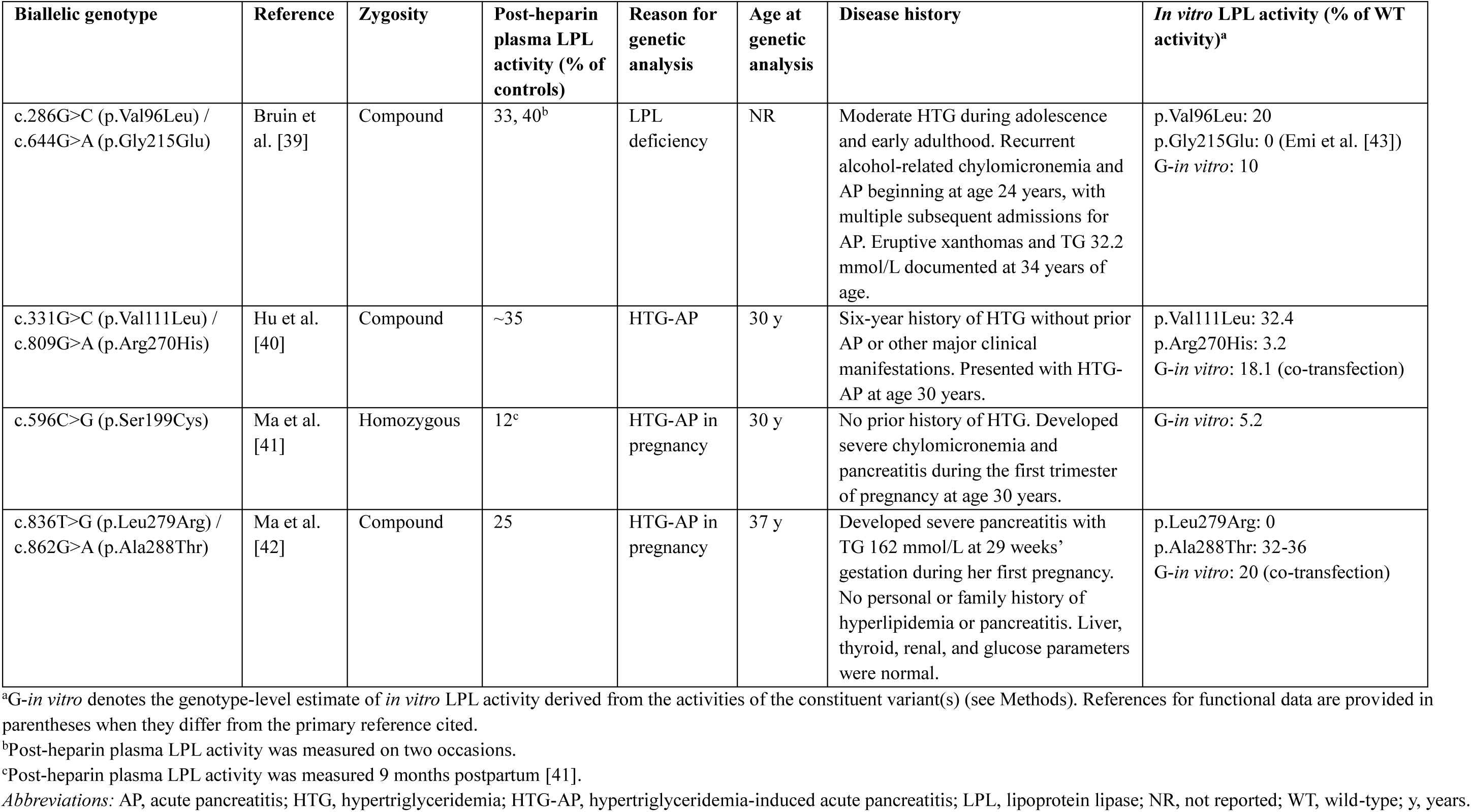
Group IV biallelic *LPL* genotypes with concordantly elevated residual LPL activity.

Of these, the homozygous c.596C>G (p.Ser199Cys) genotype exhibited the lowest residual *in vivo* and G-*in vitro* LPL activity values in the group and is discussed separately in Supplementary Note 2. Briefly, the genotype was classified using the measurement obtained 9 months postpartum, when pregnancy-related physiological changes would be expected to have largely resolved. The observed 5.2% G-*in vitro* activity was also clearly distinguishable from the background range of complete or near-complete LoF variants within the same study.

None of the four individuals exhibited the classical childhood presentation that predominated in Group I. Instead, clinically significant manifestations occurred predominantly during adulthood and were frequently associated with identifiable physiological or environmental precipitating factors. Specifically, two individuals developed severe chylomicronemia and pancreatitis during pregnancy, one experienced recurrent alcohol-related chylomicronemia and acute pancreatitis, and the remaining individual presented with HTG-associated acute pancreatitis at age 30 after a six-year history of hypertriglyceridemia (Table 2).

Considered together with the findings from Group III, these observations suggest a clinically meaningful transition at approximately 10% residual physiological LPL activity. At residual activity levels above approximately 10%, classical childhood-onset LPL deficiency was not observed in the present dataset, although clinically significant HTG and pancreatitis may still occur, particularly in the presence of additional physiological, metabolic, genetic, or environmental stressors. Group IV therefore establishes the third biological reference point in our analytical framework, whereas Group III establishes the second biological reference point, defining the biological transition zone between classical FCS and modifier-dependent clinical expression.

### Penetrance of heterozygous c/nc-LoF *LPL* variants for severe HTG

The analyses of the assembled biallelic *LPL* genotypes established three biological reference points linking residual physiological LPL activity to clinical consequence. To complement these physiological reference points and extend the framework to the heterozygous state, we next estimated the penetrance of heterozygous c/nc-LoF *LPL* variants for severe HTG.

Using the Bayesian approach described in Methods, penetrance was estimated using three quantities: *P*(G∣D), the proportion of severe HTG cases carrying the variant; *P*(G), the corresponding carrier frequency in the general population; and *P*(D), the population prevalence of severe HTG, fixed at 1:526 [44]. We first estimated penetrance at the variant-class level by considering all rare predicted LoF (pLoF) *LPL* variants collectively. This approach captures the aggregate contribution of pLoF variants and is consistent with recent studies estimating penetrance for groups of rare variants [45]. To derive *P*(G|D), we used data from the severe HTG cohort reported by Dron *et al*. [46], which represents one of the most comprehensively genotyped cohorts available for severe HTG. Five pLoF *LPL* alleles (c.46_47delCA [p.Gln16Glufs] ×2, c.127_128insT [p.Arg44Lysfs] ×1, c.272G>A [p.Trp91Ter] ×1, and c.1139+1G>A ×1) were identified among the 563 affected individuals of European ancestry. Given the rarity of biallelic rare genotypes in the severe HTG cohort [46], these alleles were used to approximate the corresponding number of heterozygous carriers, resulting in a *P*(G|D) of 0.89%. This approximation would, if anything, tend to overestimate rather than underestimate penetrance.

Because the number of controls in the Dron study was limited (503 individuals from the 1000 Genomes Project, among whom no pLoF *LPL* variants were identified) [46], we instead used rare pLoF *LPL* variants in the non-Finnish European population of gnomAD v4.1.1 (https://gnomad.broadinstitute.org/. Accessed 7 July 2026) to estimate *P*(G). After excluding the common protective c.1421C>G (p.Ser474Ter) allele (allele count = 117,901) and the rare c.1421C>A (p.Ser474Ter) allele (allele count = 1), 214 rare pLoF alleles remained among approximately 590,000 non-Finnish European individuals (Supplementary Table S4). Because none of these variants were observed in the homozygous state, the allele count corresponded to the number of heterozygous carriers, resulting in a *P*(G) of 0.036%. Using these parameters, the estimated penetrance of aggregate rare pLoF *LPL* variants for severe HTG was 4.65% (95% CI, 1.76–10.28%).

To determine whether the variant-class estimate was representative at the individual-variant level, we next performed an independent single-variant analysis using c.644G>A (p.Gly215Glu), the most frequent rare c/nc-LoF *LPL* variant identified in the severe HTG cohort [46] and which exhibited concordant near-complete loss of LPL activity in both *in vivo* and *in vitro* assays (see Supplementary Table S1). Seventeen p.Gly215Glu alleles were identified among the 563 affected individuals [46]. As above, these alleles were used to approximate the corresponding number of heterozygous carriers, resulting in a *P*(G|D) of 3.02%. In gnomAD v4.1.1, p.Gly215Glu was observed 514 times among 1,180,028 non-Finnish European alleles, corresponding to an allele frequency of 0.0004356. Because no homozygotes were observed, this represented 514 heterozygous carriers among 590,014 individuals, resulting in a carrier frequency, *P*(G), of 0.087%. Using these parameters, the estimated penetrance of p.Gly215Glu for severe HTG was 6.59% (95% CI, 3.98%–10.30%).

As summarized in Table 3, the two estimation approaches yielded remarkably consistent penetrance estimates. As a sensitivity analysis, we substituted the Ontario-based prevalence estimate of 1:400 [47] for the primary pooled estimate of 1:526 [44]. This increased the penetrance estimates from 4.65% and 6.59% to 6.12% and 8.67%, respectively. Importantly, all estimates remained below 10%, indicating consistently low penetrance despite variation in the assumed disease prevalence. Together with the three biological reference points established from the genotype–phenotype analyses of Groups I, III, and IV, these consistently low penetrance estimates complete the biological continuum by establishing the fourth biological reference point in our analytical framework.

**Table 3.**
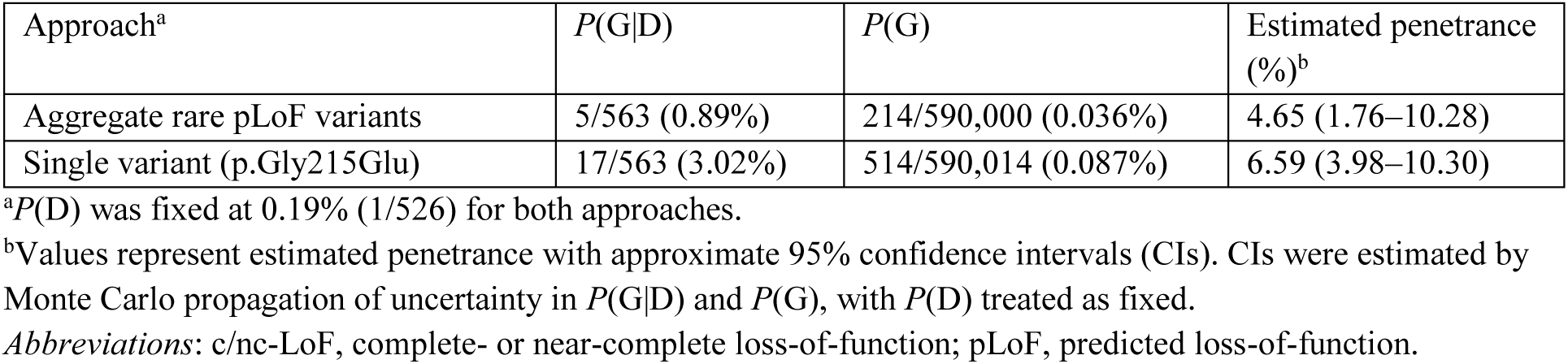
Bayesian estimates of the penetrance of heterozygous c/nc-LoF *LPL* variants for severe hypertriglyceridemia.

### Data integration

The analyses of biallelic *LPL* genotypes and the penetrance estimates for heterozygous c/nc-LoF variants establish four biological reference points that anchor the continuum of residual physiological LPL activity and clinical consequence (Fig. 2A). The first three reference points, derived from Groups I, III, and IV, delineate the relationship between residual physiological LPL activity and clinical expression, whereas the fourth provides a complementary quantitative reference point for interpreting the heterozygous state. Based on the estimated penetrance of 5–7% for severe HTG and the criteria proposed in our previous study [13], heterozygous c/nc-LoF *LPL* variants best fit the predisposing variant category.

**Figure 2.**
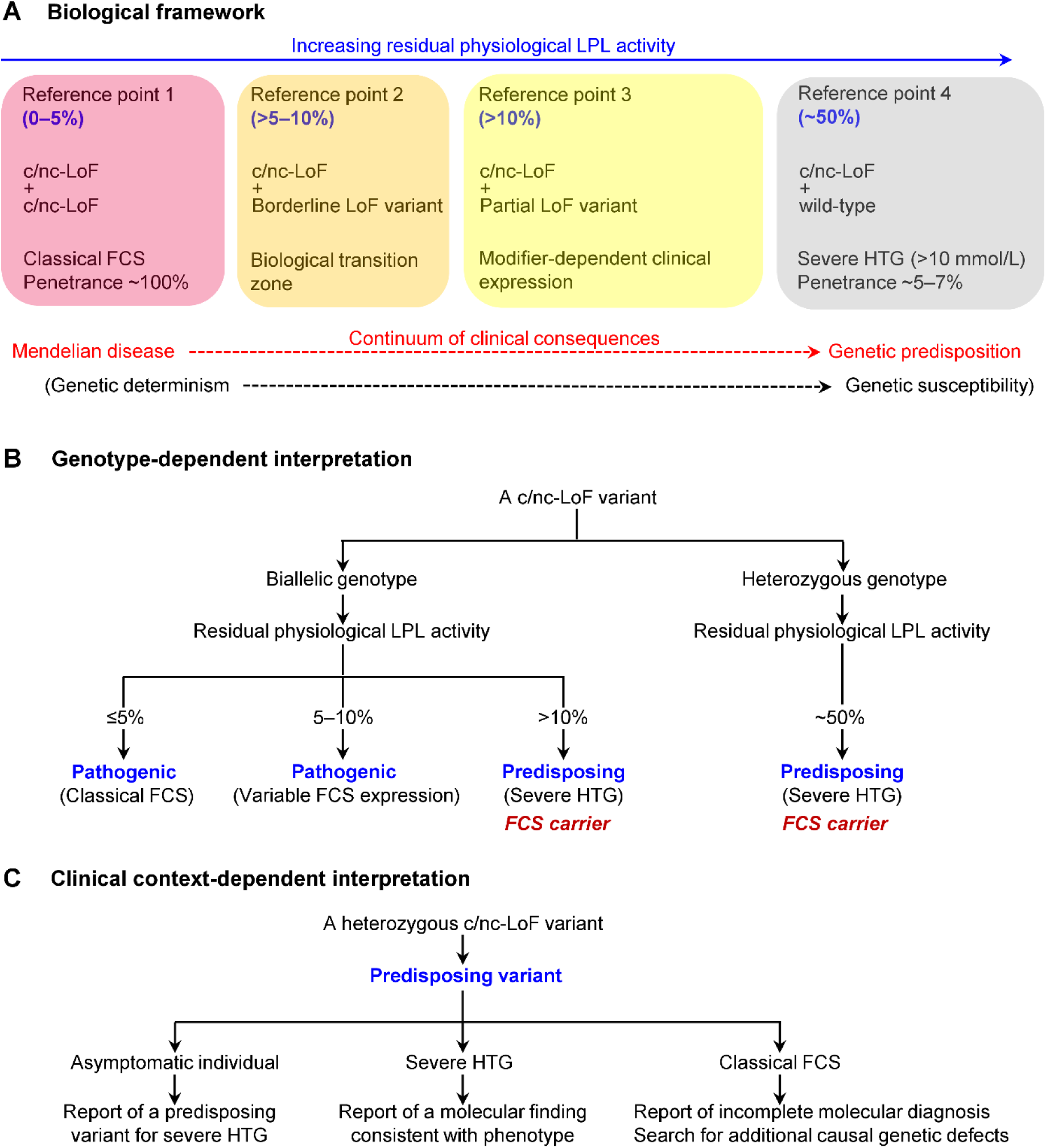
A context-dependent framework for interpreting *LPL* loss-of-function variants. (A) Illustration of the four biological reference points established in the present study, spanning the continuum from classical Mendelian disease to genetic predisposition. (B) Context-dependent interpretation framework for c/nc-LoF *LPL* variants. (C) Clinical reporting of a heterozygous c/nc-LoF *LPL* variant according to the individual’s clinical phenotype. *Abbreviations:* FCS, familial chylomicronemia syndrome; c/nc-LoF, complete- and near-complete loss-of- function; HTG, hypertriglyceridemia.

The fourth biological reference point also informs interpretation of the Group IV genotypes. Because the preceding analyses indicate a biological transition at approximately 10% residual physiological LPL activity, above which classical childhood-onset FCS becomes uncommon, severe HTG rather than FCS becomes the more appropriate clinical endpoint for considering the penetrance of Group IV genotypes. Although their extreme rarity currently precludes reliable quantitative estimation of penetrance, their substantially lower residual physiological LPL activity (12–40% of normal) than the approximately 50% expected for heterozygous c/nc-LoF variants suggests that they confer greater susceptibility to severe HTG.

Taken together, the four biological reference points define a continuum of residual physiological LPL activity underlying the transition from classical Mendelian disease to genetic predisposition.

### Developing a context-dependent framework for interpreting *LPL* LoF variants

The preceding analyses provide the basis for a biologically grounded, context-dependent framework for interpreting *LPL* LoF variants (Fig. 2B), in which residual physiological LPL activity serves as a key quantitative intermediary linking genotype to clinical consequence. Interpretation therefore depends not only on the intrinsic functional effect of an *LPL* LoF variant but also on its allelic configuration and resulting residual physiological activity. Accordingly, biallelic c/nc-LoF genotypes resulting in complete or near-complete LPL deficiency remain pathogenic for FCS, whereas heterozygous c/nc-LoF variants are more appropriately interpreted as predisposing variants for severe HTG while simultaneously indicating carrier status for FCS. Likewise, the interpretation of genotypes comprising one c/nc-LoF allele and one borderline or partial LoF allele should be guided by the residual physiological LPL activity they retain. Genotypes retaining approximately 5–10% residual physiological LPL activity may remain compatible with FCS, whereas above approximately 10% residual activity, classical FCS becomes uncommon and such genotypes are more appropriately interpreted as predisposing to severe HTG.

Fig. 2C illustrates how this framework is applied in different clinical contexts using a heterozygous c/nc-LoF *LPL* variant as an example. Although the variant classification remains unchanged, its clinical interpretation and reporting differ according to the individual’s clinical presentation. Thus, the same variant may appropriately be reported as a genetic predisposing factor in an asymptomatic individual, as a molecular finding consistent with the clinical phenotype in an individual with severe HTG, or as an incomplete molecular diagnosis requiring further investigation in an individual with classical FCS.

## Discussion

To our knowledge, this study is the first to establish a biologically grounded framework for context-dependent interpretation of *LoF* variants using *LPL* as a model gene by integrating residual physiological LPL activity measured *in vivo*, *in vitro* functional evidence, penetrance estimates, and clinical phenotype. Despite considerable methodological heterogeneity among the published *LPL* studies, four reproducible biological reference points emerged, defining a continuum from classical Mendelian disease to genetic predisposition. Together, these findings provide a biological basis for why the same *LPL* LoF variant in different allelic configurations may warrant different clinical interpretations. Several biological and clinical insights arise from this framework.

### Reconciling *in vitro* functional assays with *in vivo* physiology

As exome and genome sequencing identify increasing numbers of variants, multiplex assays of variant effect (MAVEs) are increasingly contributing to clinical variant interpretation [48, 49]. However, functional evidence should be obtained using appropriate experimental systems and interpreted within its biological context [50, 51]. Here, we observed frequent discordance between *in vitro* functional measurements and physiological LPL activity. Specifically, Group II shows that measurable residual activity in cell-based assays does not necessarily translate into physiologically meaningful residual function, whereas Group III demonstrates that complete or near-complete loss of activity *in vitro* does not preclude measurable residual physiological function *in vivo*. Finally, Group IV indicates that even when the two measurements are directionally concordant, the magnitude of residual activity may differ substantially.

Several mechanisms may explain the observed discordance between *in vitro* functional assays and physiological LPL activity. First, cell-based assays may not fully recapitulate the biological context in which LPL biosynthesis, trafficking, secretion, maturation, and function occur *in vivo*. Additional factors may include (i) unidentified variants affecting triglyceride metabolism, (ii) the simplifying assumptions used to estimate G-*in vitro* activity, and (iii) physiological and measurement variability in residual *in vivo* LPL activity. Therefore, wherever possible, functional evidence should be generated using model systems that best recapitulate the relevant biological context and interpreted in conjunction with physiological and clinical evidence to more accurately infer the biological consequences of a variant.

### Moving beyond the traditional one-variant–one-label paradigm

Earlier work recognized an uncertain boundary between familial and multifactorial chylomicronemia and proposed a continuum of increasing genetic burden from multifactorial chylomicronemia to FCS [52]. The present study extends this concept by identifying representative *LPL* genotype contexts that anchor part of this biological continuum through four reproducible biological reference points defined by residual physiological LPL activity and penetrance (Fig. 2A). Because the same c/nc-LoF variant can have fundamentally different clinical consequences depending on its biological context, context-dependent interpretation becomes necessary (Fig. 2B). This framework therefore provides a means of addressing the challenge of assigning a single “pathogenic” label to variants with context-dependent clinical consequences.

Notably, we identified approximately 10% residual LPL activity as a potential biological transition point between classical FCS and more context-dependent manifestations of severe HTG. Interestingly, approximately 10% residual CFTR function has likewise been proposed as a biologically meaningful threshold separating classical cystic fibrosis from milder *CFTR*-related disease manifestations [53]. Despite this convergence, functional thresholds defining clinically meaningful transitions are likely to be gene-specific, reflecting the distinct physiological roles and disease mechanisms of different genes associated with autosomal recessive disorders.

### Strengthening the biological foundation for the predisposing category

Within our expanded variant classification framework [13], “predisposing” variants occupy a distinct conceptual space between classical Mendelian “pathogenic” variants [5] and the recently proposed “risk” category [11], reflecting substantial functional effects with context-dependent clinical consequences. The present study supports this expanded framework by providing empirical physiological and penetrance data in the context of *LPL*. Notably, the estimated 5–7% penetrance of heterozygous c/nc-LoF *LPL* variants for severe HTG, their substantial functional effects, and their reproducible association with disease susceptibility collectively align these variants with the “predisposing” category and are consistent with their recent characterization as conferring a susceptibility state [22].

One may nevertheless argue for retaining the pathogenic designation for heterozygous c/nc-LoF *LPL* variants while explicitly acknowledging their low penetrance. Applying the pathogenic designation to heterozygous carriers may, however, imply a degree of clinical determinism that is difficult to reconcile with both their estimated penetrance and their underlying biology, particularly when more than 90% of carriers are estimated not to develop severe HTG, let alone FCS, which requires a biallelic genotype. By contrast, the predisposing designation communicates the expected clinical consequence of the heterozygous state. Such a designation represents more than a terminological refinement; it reflects the biological reality that clinically important variants may produce context-dependent spectra of clinical consequences rather than fitting neatly within conventional distinctions between Mendelian pathogenicity and benign variation [54]. Incorporating this category into the ACMG/AMP framework could therefore provide a more biologically coherent and clinically informative approach to variant interpretation in the era of precision medicine [55].

### Extending the context-dependent variant interpretation framework beyond *LPL*

Although the present study focused on *LPL*, the underlying interpretive principle is unlikely to be unique to this gene. Different genes have distinct dosage constraints, functional tolerance thresholds, and disease mechanisms. Allelic configuration may be particularly informative in recessive disorders, while in autosomal dominant disorders factors such as disease mechanism, residual biological function, penetrance, age-dependence, and genetic or environmental modifiers may assume greater relative importance. Thus, although the relevant contextual factors and resulting clinical interpretations will vary among gene–disease systems, the underlying interpretive principle may extend broadly across human disease.

Several well-established gene–disease systems illustrate this concept. Among recessive disease genes, biallelic *CFTR* variants cause cystic fibrosis, whereas certain heterozygous variants reproducibly increase susceptibility to *CFTR*-related disorders, including chronic pancreatitis [56, 57]. Likewise, biallelic *GBA1* variants cause Gaucher disease, whereas specific heterozygous variants increase susceptibility to Parkinson disease [58]. Among autosomal dominant disease genes, hypomorphic *PKD1* alleles demonstrate that different levels of residual gene function can produce distinct clinical consequences, ranging from attenuated or absent phenotypes in the heterozygous state to severe disease when inherited biallelically or in trans with another pathologically relevant allele [59].

Together with recent large-scale human genetic studies demonstrating clinically meaningful phenotypes in both monoallelic and biallelic states [14, 15], these examples suggest that, in some gene–disease systems, dominance, recessivity, Mendelian disease, and genetic predisposition may be better viewed as different regions along gene-specific biological continua rather than universally discrete categories. Accordingly, context-dependent variant interpretation may provide a broader conceptual framework in which clinical significance is shaped not by a variant’s intrinsic molecular consequence alone, but by its interaction with the biological context in which it occurs.

Incorporating biological context inevitably adds complexity, but this reflects the diversity of gene– disease relationships. Because biological function may operate along a continuum while clinical interpretation requires standardized decision thresholds, context-dependent interpretation should complement rather than replace existing classification systems. A key challenge will be to define, for individual gene–disease pairs, which contextual factors are sufficiently established to meaningfully modify variant interpretation or reporting.

### Limitations

The present study has several limitations. First, despite assembling, to our knowledge, the most comprehensive dataset of biallelic *LPL* genotypes with paired *in vivo* and *in vitro* functional data, the number of informative genotypes remained limited, particularly for Group IV. Consequently, the proposed activity ranges should be regarded as approximate biological reference points rather than precise clinical thresholds. Second, the penetrance of heterozygous c/nc-LoF *LPL* variants was estimated indirectly from published population and case-series data and is therefore subject to potential ascertainment bias, although this is unlikely to alter the principal conclusion that these variants confer a low-penetrance predisposition to severe HTG. Third, to establish proof of concept, the present study focused on the functional spectrum represented by the assembled *LPL* genotypes together with the theoretical ∼50% residual physiological LPL activity expected in heterozygous c/nc- LoF carriers. *LPL* variants with substantially milder functional effects than heterozygous c/nc-LoF variants also exist [22, 25] but were not considered here. Their biological interpretation may fall within the risk category [11, 13], illustrating that the present framework does not encompass the full *LPL* variant-effect spectrum. Delineating the biological boundary between predisposing and risk variants represents an important direction for future studies.

### Conclusions

In conclusion, the present study establishes a biologically grounded, context-dependent framework for interpreting *LPL* LoF variants by integrating physiological, functional, clinical, and population-genetic evidence. By identifying reproducible biological reference points linking residual physiological LPL activity to penetrance and clinical consequence, our analyses demonstrate that the same c/nc-LoF variant warrants different clinical interpretations depending on its allelic configuration and biological context. More fundamentally, these findings illustrate that clinical variant interpretation should integrate a variant’s molecular consequence with the biological context in which it occurs.

Although developed using *LPL* as a proof-of-concept model, the underlying interpretive principle may extend to other gene–disease systems in which clinical consequence similarly depends on biological context. As genome-first medicine expands, clinical variant interpretation may increasingly move beyond the traditional one-variant–one-label paradigm toward communicating the expected clinical consequence of a variant within its specific genotypic context. Our study thus provides a conceptual model for interpreting variants across the continuum from Mendelian disease to genetic predisposition in the era of precision medicine.

## Methods

### Study rationale and analytic strategy

In the context of LPL deficiency, complete or near-complete loss of LPL function causes FCS, whereas heterozygous c/nc-LoF variants, which are expected to reduce physiological LPL function by approximately 50%, confer increased susceptibility to severe HTG [20–22, 25, 60]. Although smaller reductions in LPL activity have also been shown to modulate TG levels [25, 61], the present study focused on the functional spectrum spanning complete LPL deficiency to the approximately 50% deficiency expected in heterozygous complete-LoF carriers, corresponding to 0% to 50% residual physiological LPL activity, as defined by biallelic complete-LoF genotypes at one end and heterozygous complete-LoF variants at the other.

To characterize the relationship between this functional spectrum and clinical phenotype, we first sought to assemble and analyze published biallelic *LPL* genotypes containing at least one missense variant. Missense variants were particularly informative because their functional consequences can range from minimal impairment to complete loss of function, thereby enabling assessment of intermediate levels of residual LPL activity. Moreover, missense variants have undergone functional characterization more frequently than most other classes of *LPL* variants [25, 62, 63]. To maximize data reliability, only biallelic genotypes for which both *in vivo* and *in vitro* LPL activity measurements were available in the same study were retained for final analysis. We reasoned that integrating genotype information with *in vivo* and *in vitro* LPL activity data and clinical phenotypes would characterize the biological continuum linking *LPL* genotype, residual physiological LPL activity, and clinical consequence. To complement the analysis of biallelic genotypes, we next estimated the penetrance of heterozygous c/nc-LoF *LPL* variants for severe HTG. Together, these complementary analyses were designed to establish the biological and quantitative basis for a conceptual framework for interpreting *LPL* LoF variants (Fig. 1).

### Reference sequences and variant nomenclature

GenBank NG_008855.2 and NM_000237.3 were used as the reference genomic and mRNA sequences for *LPL*, respectively. Individual variant nomenclature followed Human Genome Variation Society (HGVS) recommendations (https://hgvs-nomenclature.org/stable/). Notably, the nomenclature of *LPL* missense variants reported in older studies was based on the mature LPL protein sequence (448 amino acids) rather than the full-length precursor protein sequence (475 amino acids). Accordingly, all such variant designations were converted to HGVS-compliant nomenclature based on the precursor protein sequence.

### Identification and selection of biallelic genotypes

We performed a comprehensive PubMed search using the terms “LPL” or “lipoprotein lipase” in combination with “homozygous missense,” “homozygote missense,” and “compound missense” (last updated June 8, 2026). Retrieved publications were reviewed manually, and additional studies were identified through examination of cited references and a recent comprehensive review [25].

Biallelic *LPL* genotypes were included if they met all three of the following criteria: (i) the genotype contained at least one rare missense variant; (ii) *in vivo* LPL activity data (post-heparin plasma LPL activity) were available for the corresponding individual; and (iii) *in vitro* LPL activity data (measured in the medium of transfected cells following heparin treatment) for the corresponding missense variant(s) were reported in the same study. Two otherwise eligible biallelic genotypes were excluded from the final analysis because of either concerns regarding assay reliability or insufficient data for reliable quantification of residual *in vivo* LPL activity (see Supplementary Note 3).

### Data extraction

For each included genotype, the corresponding publications were manually reviewed to extract information on *in vivo* and *in vitro* LPL activity (expressed as percentages of control and wild-type values, respectively), zygosity status, the clinical indication for genetic testing, age at disease onset and age at genetic analysis, and relevant clinical history. In rare instances, a sibling carrying the same genotype as the proband was also reported. To simplify data presentation, only data from the proband were included in subsequent analyses. When the same eligible genotype was reported in more than one study, data from the first published report were usually retained for analysis unless significant discrepancies were identified between studies.

### Defining a genotype-level *in vitro* activity measure

For the included genotypes, *in vivo* LPL activity was reported at the genotype level, whereas *in vitro* LPL activity was generally reported for individual missense variant(s). To enable direct comparison at the genotype level, a genotype-level *in vitro* activity measure (designated G-*in vitro*) was defined according to the underlying genotype structure. It should be noted that all *in vitro* LPL activity measurements were derived from assays performed on the medium of transfected cells following heparin treatment.

For homozygous missense variants, G-*in vitro* was defined as the LPL activity measured in the medium of cells transfected with the corresponding mutant expression construct.

For compound heterozygous genotypes comprising a predicted LoF variant (i.e., splice-site, frameshift, or nonsense variant) and a missense variant, the predicted LoF allele was assumed to contribute zero residual LPL activity. G-*in vitro* was therefore approximated as one-half of the experimentally determined *in vitro* activity of the missense variant.

For compound heterozygous genotypes comprising two missense variants, G-*in vitro* was approximated as the mean of the corresponding *in vitro* activity values unless co-transfection data were available, in which case the co-transfection result was used.

### Estimation of the penetrance of heterozygous c/nc-LoF *LPL* variants for severe HTG

Severe HTG was selected as the clinical endpoint for this analysis because it represents the most clinically relevant manifestation of impaired TG metabolism—in contrast to the highly prevalent mild-to-moderate HTG, which affects approximately one-quarter of North Americans [47]—and is also the hallmark biochemical phenotype of FCS. A TG threshold of >10 mmol/L (>885 mg/dL), consistent with several European guidelines [64–67], was adopted because compatible estimates of both the epidemiological prevalence of severe HTG and the proportion of affected individuals carrying c/nc- LoF *LPL* variants were available for this definition.

Penetrance was estimated using Bayes’ theorem, which integrates disease prevalence, variant frequency among affected individuals, and variant frequency in the general population to estimate the probability of disease among variant carriers [13, 45]. Specifically, let D denote severe HTG and G heterozygous carriage of the c/nc-LoF LPL variant(s) under investigation. The penetrance of heterozygous c/nc-LoF *LPL* variant(s) for severe HTG, *P*(D|G), was estimated from three epidemiological quantities: (i) the proportion of individuals with severe HTG carrying the c/nc-LoF *LPL* variant(s), *P*(G|D); (ii) the prevalence of severe HTG in the general population, *P*(D); and (iii) the frequency of heterozygous carriers of the c/nc-LoF LPL variant(s) in the general population, *P*(G). According to Bayes’ theorem,

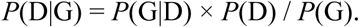

The primary estimate of *P*(D) (1:526) was obtained from a recent pooled analysis of epidemiological studies of severe HTG [44] and was used for two complementary penetrance analyses. An Ontario-based estimate (1:400) [47] was evaluated separately in sensitivity analyses.

The first analysis estimated penetrance for the aggregate class of rare predicted LoF (pLoF) *LPL* variants (variant-class analysis). This approach captures the aggregate contribution of pLoF variants and is consistent with recent studies estimating penetrance for groups of rare variants [45]. An independent single-variant analysis was then performed using c.644G>A (p.Gly215Glu) to complement the aggregate variant-class analysis. In both analyses, *P*(G|D) was derived from the severe HTG cohort reported by Dron et al. [46], which comprises 563 individuals of European ancestry and represents one of the largest and most comprehensively genotyped severe-HTG cohorts currently available. *P*(G) was estimated from the non-Finnish European population of the Genome Aggregation Database (gnomAD v4.1.1; https://gnomad.broadinstitute.org/).

Approximate 95% confidence intervals for penetrance were estimated by Monte Carlo propagation of uncertainty in *P*(G|D) and *P*(G) through the Bayes’ theorem equation while treating *P*(D) as fixed. For each analysis, 1,000,000 paired values of *P*(G|D) and *P*(G) were independently sampled from Beta(*x* + 0.5, *n − x* + 0.5) distributions derived from the corresponding observed binomial counts under Jeffreys’ prior [68]. Penetrance was calculated for each paired draw, and the 2.5th and 97.5th percentiles of the simulated penetrance distribution were taken as the approximate 95% confidence interval.

## Data availability

All data supporting this study are provided within the article and its supplementary information.

## Acknowledgments

We thank the original authors who reported the *LPL* variants analyzed in this study. This study was supported by the Institut National de la Santé et de la Recherche Médicale (INSERM), France and the National Key Research and Development Program of China (2025YFC2511900). D.N.C. wishes to acknowledge financial support from Qiagen Inc through a License Agreement with Cardiff University. The funding bodies did not play any role in the study design, collection, analysis, and interpretation of data or the writing of the article and the decision to submit it for publication.

## Author contributions

Conceptualization and design, J.-M.C., Q.Y., W.-B.Z., and W.L.; data collation, J.-M.C., Q.Y., N.P., Y.H., X.L.; data interpretation, J.-M.C., Q.Y., W.-B.Z., N.P., Y.L., Y.H., Y-C.W., X.L., E.G., E.M., J.W., C.F., D.N.C., and W.L.; writing – original draft, J.-M.C.; writing – review & editing, J.-M.C., Q.Y., W.-B.Z., N.P., Y.L., Y.H., Y-C.W., X.L., E.G., E.M., J.W., C.F., D.N.C., and W.L. All authors read and approved the final version of the manuscript and agree to be accountable for all aspects of the work, ensuring that questions related to the accuracy or integrity of any part are appropriately investigated and resolved.

## Declaration of generative AI and AI-assisted technologies in the writing process

During the preparation of this work the authors used ChatGPT Plus in order to improve readability and language. After using this tool, the authors reviewed and edited the content as needed and take full responsibility for the content of the publication.

