## Supplementary file for "Context-dependent variant interpretation from Mendelian disease to genetic predisposition: a proof-of-concept using *LPL*"

**Supplementary Table S1. Group I biallelic *LPL* genotypes with concordant complete or near-complete functional loss (N = 27)**

| **Biallelic *LPL* genotype** | **Ref.** | **Zygosity** | **Post-heparin plasma LPL activity (% of controls)** | **Reason for genetic analysis** | **Age at genetic diagnosis** | **Disease history** | ***In vitro* LPL activity (% of WT activity)[a]** |
|---|---|---|---|---|---|---|---|
| c.[106G>A;865T>C] (p.[Asp36Asn;Tyr289His]) | [1] | Hom | 0 | LPL deficiency | 62 y | Recurrent abdominal pain since childhood; diagnosed at age 22 years with lipemia retinalis, marked hepatosplenomegaly, HTG, and chylomicronemia. | p.Asp36Asn: 85<br>p.Tyr289His: 0<br>p.[Asp36Asn;Tyr289His]: 0<br>G-*in vitro*: 0 |
| c.209A>G (p.Asn70Ser) | [2] | Hom | 0 | Chylomicronemia | 20 y | Chylomicronemia diagnosed during infancy; recurrent pancreatitis. | G-*in vitro*: 0 |
| c.337T>C (p.Trp113Arg) / c.397C>T (p.Gln133*) | [3] | CH | <2 | Type I hyperlipoproteinemia | 1 y | Presented at age 11 days with jaundice, eruptive xanthomas, and lipemia retinalis. | p.Trp113Arg: 1.3<br>p.Gln133*: 0<br>G-*in vitro*: 0.66 |
| c.394G>A (p.Gly132Arg) / c.693C>G (p.Asp231Glu) | [4] | CH | 0 | Type I hyperlipoproteinemia | 21 m | Very severe HTG (28.8 mmol/L) | p.Gly132Arg: 0<br>p.Asp231Glu: 0<br>G-*in vitro*: 0 |
| c.496G>A (p.Gly166Ser) | [5] | Hom | 0 | LPL deficiency | 13 y | Hospitalized at age 29 days with persistent diarrhea; lipemia retinalis and milky plasma were noted. | G-*in vitro*: 0 |
| c.542G>T (p.Gly181Val) / c.1322+2T>C | [6] | CH | 0 | Type I hyperlipoproteinemia | 2 m | Severe HTG (22.39 mmol/L) | p.Gly181Val: 0<br>c.1322+2T>C: 0<br>G-*in vitro*: 0 |
| c.[548A>G;1421C>G] (p.[Asp183Gly;Ser474*]) | [7] | Hom | 0 | FCS | 12 y | Acute pancreatitis, marked hepatosplenomegaly, and lipemia retinalis at age of 1 year. | p.[Asp183Gly;S474*]: 0<br>G-*in vitro*: 0 |
| c.548A>G (p.Asp183Gly) | [8] | Hom | 0 | LPL deficiency | 18 y | Presented with abdominal pain resulting in laparotomy at age 17 years. | G-*in vitro*: 0 |
| c.547G>A (p.Asp183Asn) / c.727T>A (p.Cys243Ser) | [8] | CH | BD | LPL deficiency | 38 y | Presented at age 25 years with recurrent episodes of pancreatitis and chylomicronemia. | p.Asp183Asn: 0<br>p.Cys243Ser: 0<br>G-*in vitro*: 0 |
| c.551C>G (p.Pro184Arg) | [9] | Hom | 1.4 | Type I hyperlipoproteinemia | 32 y | Presented at age 6 weeks with chyloperitoneum requiring laparotomy; milky serum, hepatosplenomegaly, and chylothorax were noted. | G-*in vitro*: 0 |
| c.607G>A (p.Ala203Thr) | [10] | Hom | 2.8 | LPL deficiency | 39 y | Presented at age 21 years with recurrent abdominal pain and pancreatitis since childhood; evaluation revealed lipemia retinalis and splenomegaly. | G-*in vitro*: 2.6 |
| c.621C>G (p.Asp207Glu) | [11] | Hom | 4 | FCS | 10 y | Diagnosed based on marked fasting hypertriglyceridemia, chylomicronemia, and abdominal pain. | G-*in vitro*: 0 |
| c.628C>G (p.His210Asp) / c.89-4_-2del | [12] | CH | <1 | Primary hyperchylomicronemia | 5 y | Presented with hepatomegaly, splenomegaly and a milky fasting serum. | p.His210Asn: 0<br>c.89-2_-4del: 0<br>G-*in vitro*: 0 |
| c.644G>A (p.Gly215Glu) | [13] | Hom | BD | LPL deficiency | 29 y | Since early infancy, had recurrent episodes of acute abdominal pain. Lipid analysis revealed severe hypertriglyceridemia (1,500–2,500 mg/dL) and fasting chylomicronemia. | G-*in vitro*: <1 |
| c.662T>C (p.Ile221Thr) | [14] | Hom | 0 | LPL deficiency | 4 y | Presented at age 4 years with hepatosplenomegaly and abdominal pain. Plasma lipid analysis revealed chylomicronemia with triglyceride levels exceeding 30 mmol/L. | G-*in vitro*: 0 |

| | | | | | | | |
|---|---|---|---|---|---|---|---|
| c.662T>C (p.Ile221Thr) / c.809G>A (p.Arg270His) | [15] | CH | <1 | LPL deficiency | 34 y | Throughout childhood, experienced recurrent episodes of abdominal pain and pancreatitis, accompanied by eruptive xanthomas, lipemia retinalis, and splenomegaly. | p.Ile221Thr: 1.4<br>p.Arg270His: <1<br>G-*in vitro*: <1.2 |
| c.665G>A (p.Gly222Glu) | [16] | Hom | 0 | LPL deficiency | 9 y | At age 3 years, developed acute pancreatitis with plasma triglyceride levels exceeding 7,000 mg/dL (79 mmol/L), hepatomegaly, and eruptive xanthomas. | G-*in vitro*: 0 |
| c.680T>C (p.Val227Ala) / c.1139+1G>A | [17] | CH | 0 | Excessive primary HTG | 37 y | At age 37 years, was admitted for severe primary hypertriglyceridemia. Medical history revealed recurrent pancreatitis and triglyceride levels >2,000 mg/dL. | p.Val227Ala: 0<br>c.1139+1G>A: 0<br>G-*in vitro*: 0 |
| c.701C>T (p.Pro234Leu) / c.808C>T (p.Arg270Cys) | [18] | CH | 0 | LPL deficiency | At birth | Presented with chylomicronemia at birth, eruptive xanthomas at age 3 years, pancreatitis at age 9 years, and hepatomegaly. | p.Pro234Leu: 0 [19]<br>p.Arg270Cys: 0<br>G-*in vitro*: 0 |
| c.755T>C (p.Ile252Thr) / c.809G>A (p.Arg270His) | [20] | CH | 0 | LPL deficiency | NR | Presented in early childhood with classical LPL deficiency, including acute pancreatitis, chylomicronemia (triglyceride, 25 mmol/L), and complete absence of post-heparin plasma LPL activity. | p.Ile252Thr: ~10<br>p.Arg270His: 0 [21]<br>G-*in vitro*: ~5 |
| c.811T>A (p.Ser271Thr) / c.250-1G>A | [22] | CH | 0 | LPL deficiency | 6 y | A 6-year-old boy who had experienced recurrent episodes of abdominal pain and eruptive xanthomas since early infancy, particularly during febrile illnesses. Lipid analysis revealed severe hypertriglyceridemia (1,900–4,800 mg/dL) and fasting chylomicronemia. | p.Ser271Thr: 0<br>c.250-1G>A: 0<br>G-*in vitro*: 0 |
| c.829G>A (p.Asp277Asn) / c.833C>G (p.Ser278Cys) | [23] | CH | 0 | LPL deficiency | 16 y | At age 2 years, was incidentally found to have milky plasma and was subsequently diagnosed with LPL deficiency based on absent post-heparin plasma LPL activity. | p.Asp277Asn: 3 [24]<br>p.Ser278Cys: 0<br>G-*in vitro*: 1.5 |
| c.858T>A (p.Ser286Arg) | [25] | Hom | 0 | LPL deficiency | 6 | At age 6 months, asymptomatic chylomicronemia was detected. At age 18 months, abdominal pain developed despite a normal diet. Examination revealed hepatosplenomegaly, eruptive xanthomas, lipemia retinalis, and severe hypertriglyceridemia (2,900 mg/dL). | G-*in vitro*: 2 |
| c.865T>C (p.Tyr289His) | [1] | Hom | 0 | LPL deficiency | 62 y | Recurrent abdominal pain since childhood. Evaluation at age 22 years revealed lipemia retinalis, marked hepatosplenomegaly, hypertriglyceridemia, and chylomicronemia. | G-*in vitro*: 0 |
| c.891T>G (p.Phe297Leu) | [26] | Hom | 0 | Milky appearance of the fasting plasma | 1 m | Presented with eruptive xanthomas and lipemia retinalis. | G-*in vitro*: 0 |
| c.1211T>G (p.Met404Arg) | [27] | Hom | 0 | Type I hyperlipoproteinemia | 28 y | Selected for genetic analysis based on fasting triglyceride levels of 10 mmol/L, at least one episode of pancreatitis, absence of secondary causes of hypertriglyceridemia, and family consanguinity. No further clinical details were available. | G-*in vitro*: 0 |
| c.[1310A>T;1421C>G] (p.[Glu437Val;Ser474*]) | [28] | Hom | 0 | LPL deficiency | 3 y | Presented at age 2 months with acute pancreatitis and hepatosplenomegaly. Plasma lipid analysis revealed chylomicronemia with triglyceride levels exceeding 12,000 mg/dL. | G-*in vitro*: 0 |

The earliest reported age, defined as either age at symptom onset or, when unavailable, age at genetic diagnosis, is highlighted in red when ≤10 years. Childhood was operationally defined as ≤10 years of age. Of the 27 biallelic genotypes, 23 met this criterion, whereas the remaining four genotypes are highlighted in yellow.

[a]G-*in vitro* denotes the genotype-level estimate of *in vitro* LPL activity derived from the activities of the constituent variant(s) (see Methods). References for functional data are provided in parentheses where they differ from the primary reference cited.

*Abbreviations:* BD, barely detectable; CH, compound heterozygote; FCS, familial chylomicronemia syndrome; Hom, homozygote; HTG, hypertriglyceridemia; LPL, lipoprotein lipase; NR, not reported; WT, wild-type; y, years.

**Supplementary Table S2. Group II biallelic *LPL* genotypes with discordantly elevated G-*in vitro* LPL activity**

| **Biallelic genotype** | **Reference** | **Post-heparin plasma LPL activity (% of controls)** | **Reason for genetic analysis** | **Age at genetic analysis** | **Disease history** | ***In vitro* LPL activity (% of WT activity)[a]** |
|---|---|---|---|---|---|---|
| c.382A>G (p.Thr128Ala) / c.829G>A (p.Asp277Asn) | [29] | <0.9 | Chylomicronemia | 62 y | History of recurrent abdominal pain since childhood. First evaluated at age 39 following an episode of acute pancreatitis (triglyceride, 36.1 mmol/L). | p.Thr128Ala: 27.6<br>p.Asp277Asn: 3 (Ishimura-Oka et al. [24])<br>G-*in vitro*: 15.3 |
| c.662T>C (p.Ile221Thr) / c.1334G>A (p.Cys445Tyr) | [30] | 0 | LPL deficiency | 30 y | Diagnosed with LPL deficiency in childhood. | p.Ile221Thr: 0 (Henderson et al. [14]; Dichek et al. [15])<br>p.Cys445Tyr: 48<br>G-*in vitro*: 25 (co-transfection) |
| c.953A>G (p.Asn318Ser) / c.1019-3C>A | [17] | 0 | Severe HTG | 40 y | Severe HTG (39.5 mmol/L; 3500 mg/dL) was first documented at 18 years of age and decreased to <5.6 mmol/L (<500 mg/dL) under a strict diet. No history of acute pancreatitis was reported. | p.Asn318Ser: 50–70 (Reymer et al. [31]; Zhang et al. [32]; Busca et al. [33]; Syvanne et al. [34])[b]<br>c.1019-3C>A: 0<br>G-*in vitro*: 25–35 |

[a]G-*in vitro* denotes the genotype-level estimate of *in vitro* LPL activity derived from the activities of the constituent variant(s) (see Methods). References for functional data are provided in parentheses where they differ from the primary reference cited.
*Abbreviations:* HTG, hypertriglyceridemia; LPL, lipoprotein lipase; WT, wild-type; y, years.

**Supplementary Table S3. Biallelic *LPL* genotypes with discordantly elevated residual *in vivo* LPL activity excluded from the final Group III analysis**

| Biallelic genotype | Reference | Zygosity | Post-heparin plasma LPL (% of controls) | Reason for genetic analysis | Age at genetic analysis | Disease history | *In vitro* LPL activity (% of WT activity)[a] |
|---|---|---|---|---|---|---|---|
| c.693C>G (p.Asp231Glu) | [21] | Homozygous | 19 | HTG | 44 y | Asymptomatic; diagnosed incidentally at age 42 years during evaluation for HTG. No eruptive xanthomas, lipemia retinalis, pancreatitis, or hepatosplenomegaly were reported. | G-*in vitro*: 0 |
| | [35] | Homozygous | 27.6 | Severe HTG and recurrent pancreatitis | 39 y | Presented with recurrent pancreatitis and severe HTG (serum TG 2032 mg/dL [≈23.0 mmol/L]). No hepatosplenomegaly, eruptive xanthomas, or cardiovascular disease were reported. No obesity, diabetes mellitus, or heavy alcohol intake was documented. | G-*in vitro*: 0 |
| c.809G>A (p.Arg270His) | [21] | Homozygous | 17 | LPL deficiency | 23 d | Presented with eruptive xanthomas, lipemia retinalis, and hepatosplenomegaly. | G-*in vitro*: 0 |

See Supplementary Note 1 for why they were excluded from final analysis.

[a]G-*in vitro* denotes the genotype-level estimate of *in vitro* LPL activity derived from the activities of the constituent variant(s) (see Methods).

**Supplementary Note 1. Biallelic *LPL* genotypes with discordantly elevated residual *in vivo* LPL activity excluded from the final Group III analysis**

Among the initial Group III genotypes, three individuals representing two homozygous genotypes (c.693C>G [p.Asp231Glu], n = 2; c.809G>A [p.Arg270His], n = 1) exhibited markedly elevated residual *in vivo* activities (17%–27.6%) despite complete absence of G-*in vitro* activity (Supplementary Table S3). The highest *in vivo* LPL activities, 19% and 27.6% of control values, were reported in two individuals homozygous for c.693C>G (p.Asp231Glu) [21, 35]. These findings were generally consistent with the corresponding clinical phenotypes, particularly in the individual reported by Gotoda et al. [21], who remained asymptomatic until age 42 years, when severe HTG was diagnosed incidentally. However, in the Gotoda study [21], the homozygous c.693C>G (p.Asp231Glu) genotype was reported alongside the homozygous c.809G>A (p.Arg270His) genotype, which had the third-highest *in vivo* LPL activity (17%), as well as three homozygous nonsense or splice-site genotypes. Notably, the absolute post-heparin plasma LPL activities were remarkably similar across all five homozygous genotypes, ranging from 0.8 to 1.2 µmol FFA/mL/h (control mean, 6.4 ± 2.1 µmol FFA/mL/h; FFA, free fatty acids) [21]. Thus, the reported percentages for the homozygous c.693C>G (p.Asp231Glu) and c.809G>A (p.Arg270His) genotypes may overestimate the extent of physiologically relevant residual LPL activity. Indeed, by comparison with the three homozygous nonsense or splice-site genotypes in the same study, which exhibited similar absolute post-heparin plasma LPL activities, the reported 19% value for the homozygous c.693C>G (p.Asp231Glu) genotype and 17% value for the homozygous c.809G>A (p.Arg270His) genotype may correspond to trace residual LPL activity. This interpretation would be consistent with both the absence of detectable G-*in vitro* activity (0%) and the classical FCS phenotype associated with the homozygous c.809G>A (p.Arg270His) genotype, as evidenced by the affected individual's presentation at 23 days of age with eruptive xanthomas, lipemia retinalis and hepatosplenomegaly (Supplementary Table S3).

However, this interpretation is less straightforward for the homozygous c.693C>G (p.Asp231Glu) genotype, given both the mild phenotype reported by Gotoda et al. [21] and the independently reported *in vivo* activity of 27.6% in a second homozygous individual [35]. The latter individual had severe HTG and recurrent pancreatitis but lacked several classical FCS manifestations, including hepatosplenomegaly and eruptive xanthomas, and had no documented obesity, diabetes mellitus, or heavy alcohol intake [35]. Although the basis for these observations remains uncertain, Murano et al. [35] showed that p.Asp231Glu LPL had markedly reduced activity toward very low-density lipoproteins and conventional triolein substrates but exhibited restored activity when triolein was emulsified with acidic phospholipids such as phosphatidylethanolamine, phosphatidylserine, or cardiolipin. These findings suggest that the functional consequences of p.Asp231Glu depend on substrate or lipoprotein-surface composition and, consequently, on the physiological lipoprotein environment.

Taken together, these observations indicate that the reported *in vivo* activities for these three individuals require cautious interpretation and should not be interpreted as representative of the biological transition zone defined by the remaining Group III genotypes.

**Supplementary Note 2. The homozygous c.596C>G (p.Ser199Cys) genotype in Group IV**

This genotype was identified in a 30-year-old female who developed HTG-associated acute pancreatitis during pregnancy despite having no prior history of HTG [36]. Post-heparin plasma LPL activity was measured at four time points after delivery (10 days, 6 weeks, 4 months, and 9 months postpartum), yielding values of 0, 5.0, 46.7, and 33.3 nmol FFA/min/mL, respectively. The 12% value used in the present study was derived from the measurement obtained 9 months postpartum, when physiological changes associated with pregnancy would be expected to have largely resolved.

Eight additional missense variants analyzed in the same study all exhibited activities of 0–1% of wild-type activity [36]. Thus, the observed G-*in vitro* activity of 5.2% for p.Ser199Cys was clearly distinguishable from the range associated with complete or near-complete loss of function.

**Supplementary Note 3. Rationale for excluding two biallelic *LPL* genotypes from the study dataset**

Two otherwise eligible genotypes were excluded from the final analysis. The first was the compound heterozygous genotype c.188C>T (p.Ser63Phe) and c.662T>C (p.Ile221Thr) [37]. It was excluded because of concerns regarding the reliability of the conditioned-medium LPL activity assay. Specifically, the empty-vector control exhibited approximately one-third of the activity of the wild-type expression construct, whereas another variant analyzed in the same study, c.835_836del (p.Leu279Valfs*3), displayed LPL activity comparable to that of the wild-type construct [37]. The second excluded genotype was the homozygous c.835C>G (p.Leu279Val) genotype [38]. The published study did not provide sufficiently clear genotype-specific clinical and functional data to permit reliable integration into the present genotype-level analysis [38]. Consequently, this genotype was excluded from the final dataset.
